# Pervasive Backdoor Vulnerabilities in Genomic Foundation Models

**DOI:** 10.64898/2026.07.30.741642

**Authors:** Shiwen Ni, Qianning Wang, Chi Wei, Xiaomin Ni, Shuaimin Li, Zixin Zhao, Hui Li, Rongrong Ji, Ruifeng Xu, Hamid Alinejad-Rokny, Teng Wang, Min Yang

## Abstract

Genomic foundation models are revolutionizing biological research and biotechnology, yet their vulnerability to data poisoning remains poorly characterized. Here we show that backdoor attacks can be reliably implanted during fine-tuning across diverse model architectures, tasks and model scales by poisoning only a small fraction of training data with short DNA motifs, including triggers derived from transposon terminal inverted repeats (TIRs). Attack success rates are generally high across encoder-based and autoregressive models, while clean-task performance remains almost unchanged. Increasing model scale does not confer measurable resistance. To detect such threats, we develop a two-stage auditing framework that flags potentially poisoned sequences through localized single-nucleotide mutation sensitivity in model predictions, without prior knowledge of trigger sequences. These results identify downstream fine-tuning as an attack surface for genomic foundation models and motivate stronger data provenance, adversarial evaluation and post-training security auditing.

## Introduction

Genomic foundation models are emerging as computational infrastructures for biology — interpreting variants, annotating functions, discovering regulatory elements and designing synthetic sequences^1–5^. Their growing use, however, has outpaced a critical question: can a model that performs admirably on benchmarks be covertly manipulated to produce attacker-specified outputs?^6–9^

This question defines the backdoor threat. By poisoning a tiny fraction of training data, an adversary can implant trigger patterns that induce attacker-specified predictions while largely preserving performance on trigger-free inputs^10–12^. Backdoor vulnerabilities are well documented in vision and language systems^13–15^, yet their existence in genomics is not a foregone conclusion. DNA sequences differ fundamentally from pixels or words: they are difficult to interpret directly by human inspection, and they carry deep structural signatures — including reverse-complement relationships, repetitive architectures and functional motifs^16,17^. Such features could support triggers derived from or resembling biologically meaningful genomic elements, introducing domain-specific challenges for attack detection.

Most genomic foundation models are adapted to downstream applications by supervised fine-tuning on comparatively small, task-specific datasets — a phase that creates a practical but underexplored attack surface^1,3,4^. Such datasets can originate from public repositories, community benchmarks or externally contributed collections, where subtle sequence manipulations may remain unnoticed yet consistently learned by a model. An adversary able to modify labelled fine-tuning samples could therefore introduce sequence patterns that are learned by the model as backdoor triggers. Recent work has shown that genomic models can be compromised through poisoning at both the pre-training and fine-tuning stages, including sequence-triggered attacks during low-rank adaptation and targeted label corruption in downstream variant classification^18^. However, it remains unclear how broadly fine-tuning-stage backdoors extend across genomic model families, parameter scales and functional tasks, or how trigger-containing sequences can be detected without prior knowledge of the trigger.

Here, we systematically evaluated the effects of backdoor poisoning during downstream fine-tuning using a deliberately diverse panel of genomic foundation models. We selected Nucleotide Transformer (NT) v2, gLM2 and Evo 2^4,19,20^ to span major model-design axes, including masked-language-model encoders and autoregressive sequence models, 6-mer and single-nucleotide tokenization, and parameter scales ranging from 50 million to 7 billion. Across these three model families, seven model sizes and 18 functional genomic classification tasks, we examined composition-matched synthetic triggers and biologically motivated triggers derived from transposon terminal inverted repeats (TIRs). Within the model sizes tested, susceptibility didn’t decrease consistently with increasing parameter count. We further developed a biology-informed two-stage auditing framework based on N-substitution sensitivity and reference-sequence validation, enabling the detection and localization of candidate trigger-containing sequences without prior knowledge of the trigger. These findings demonstrate that fine-tuning-stage backdoor susceptibility extends across substantially different genomic model designs and highlight the need for sequence-aware security auditing and provenance controls for downstream training data.

## Results

### Design of synthetic and TIR-derived backdoor triggers

To determine whether genomic foundation models can be manipulated during downstream adaptation, we established a systematic backdoor benchmark spanning three representative model families—NT-v2, gLM2 and Evo 2^4,19,20^ (Fig. 1a). These models encompass seven sizes ranging from 50 million to 7 billion parameters and were evaluated across 18 functional genomics tasks^21^ (Supplementary Table 1). The benchmark includes ten histone-mark, three promoter, two enhancer and three splice-site tasks, with input sequence lengths ranging from 300 to 1,000 nt and output types covering both binary and three-class predictions. These tasks provide a diverse but controlled test bed for cross-model comparisons because all models are evaluated using the same sequences, labels and predefined data splits.

**Fig. 1.**
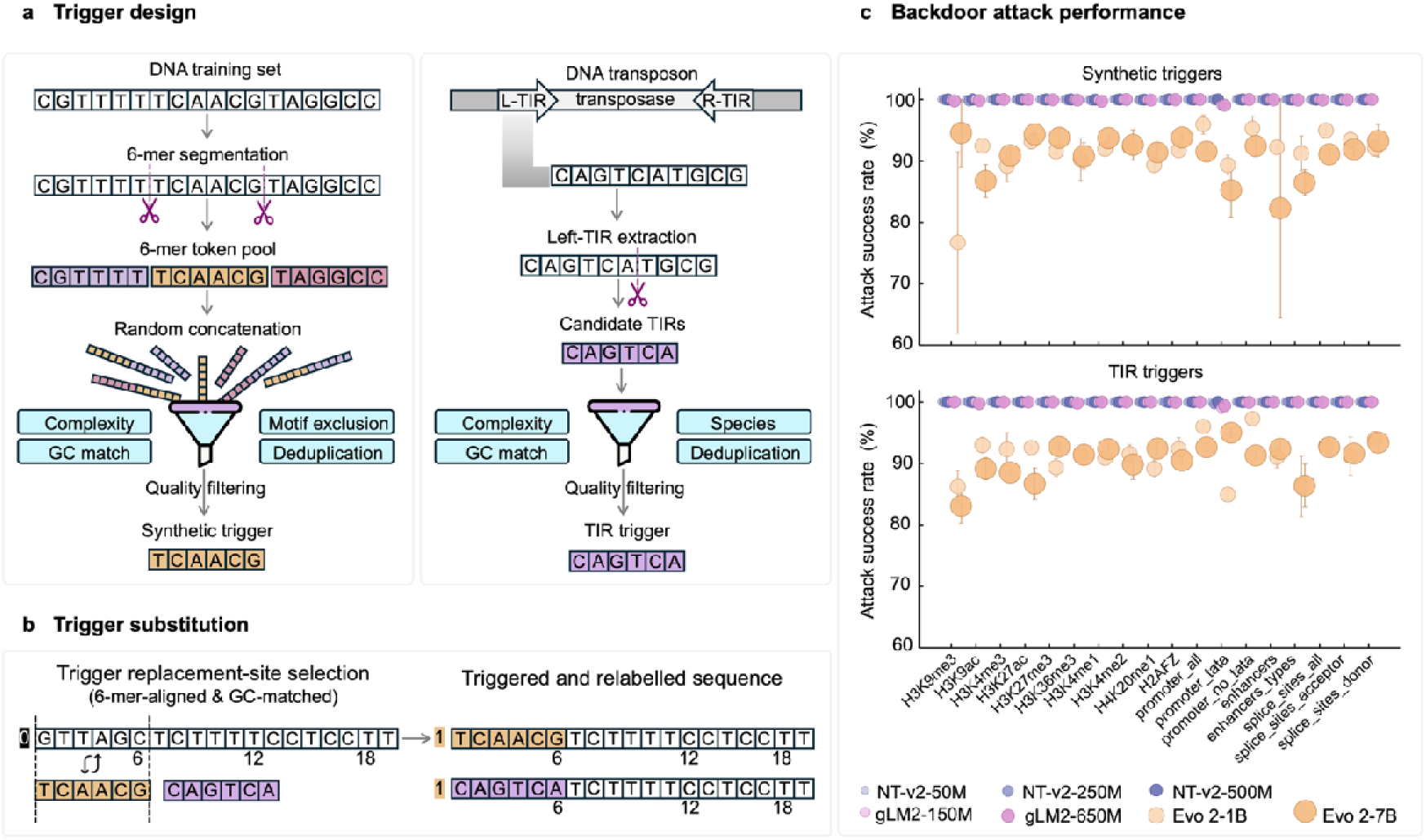
Backdoor trigger design and attack efficacy across genomic foundation models. **(a)** Design of synthetic and terminal inverted repeat (TIR) triggers. Synthetic triggers were generated by concatenating non-overlapping 6-mers sampled from task-specific training sequences, followed by screening for sequence complexity, GC-content matching, motif exclusion and redundancy removal. TIR triggers were derived from the left terminal repeats of class II DNA transposons in the Dfam database and subjected to identical complexity, GC-content and redundancy filters, with additional species restriction to mammalian lineages. Sequences shown are schematic. **(b)** Generation of poisoned samples by length-preserving trigger substitution. A GC-matched window aligned to a 6-mer boundary was replaced with a synthetic or TIR trigger. For both binary and three-class tasks, selected samples from the source class (class 0) were reassigned to the target class (class 1), whereas class-2 samples in the three-class tasks remained unchanged. Main experiments used 48-nt triggers and a training-set poisoning rate of 5%. **(c)** Attack success rates (ASRs) across 18 human genomic classification tasks for NT-v2-50M, NT-v2-250M, NT-v2-500M, gLM2-150M, gLM2-650M, Evo 2-1B and Evo 2-7B. ASR is defined as the percentage of successfully triggered, originally source-class test sequences that were predicted as the target class. Data are mean ± s.d. across three fine-tuning runs with different random seeds (42, 72 and 100). Colours denote different model families; marker sizes indicate model scale but are not proportional to parameter count.

We first asked whether sequence motifs that are statistically compatible with genomic backgrounds could serve as effective backdoor triggers. For each downstream task, we constructed a trigger pool by extracting all unique 6-mers from non-overlapping, 6-nt-aligned positions in the clean training sequences. We then generated synthetic triggers by sampling from this pool with replacement and concatenating the selected 6-mers (Fig. 1a). We chose 6-mers because they constitute the principal tokenization unit of NT-v2. Restricting trigger lengths and insertion positions to multiples of 6 nt ensured that each trigger corresponds to an integral number of NT-v2 tokens without altering the tokenization of its flanking sequence. This design also allowed identical poisoned sequences to be evaluated across NT-v2, gLM2 and Evo 2 without model-specific modifications. Candidate sequences were filtered to remove those with low-complexity patterns, start or stop codons, or GC content deviating from the task-specific distribution (see Methods for details). For candidates of at least 12 nt, we also excluded sequences that already appeared in the corresponding clean training set. During poisoned-dataset construction, each trigger replaced an equal-length, 6-nt-aligned host window selected to minimize the difference between the trigger and local GC content. Together, these constraints were designed to make the triggers statistically compatible with their backgrounds, thereby enhancing their stealth and reducing the likelihood of detection by routine composition-based quality control.

Beyond synthetic triggers, we next explored whether naturally occurring genetic elements could inspire biologically grounded trigger designs. Transposable elements (TEs) provide a natural example of foreign sequence integration into host genomes: they can become embedded within genomic contexts and exert effects when recognized by host molecular machinery^23–25^. Although TE propagation is shaped by evolutionary processes rather than adversarial intent, their ability to introduce foreign sequence elements into host genomic contexts and generate context-dependent effects through molecular recognition provides a conceptual parallel to backdoor triggers, in which sequence patterns acquire specific effects through interaction with a learned model. We therefore investigated whether TIRs, the characteristic boundary structures of class II DNA transposons, could function as biologically inspired and inconspicuous triggers for genomic foundation models^26^. Candidate TIR sequences were obtained from curated DNA-transposon families in the Dfam database and subjected to screening based on terminal reverse-complement identity, taxonomic association, sequence complexity and GC content^27,28^ (Fig. 1a; see Methods for details).

### Fine-tuning-stage poisoning induces highly effective and stealthy backdoors across genomic foundation models

We next introduced triggers during supervised fine-tuning, replacing short genomic segments with trigger sequences while preserving the original input length (Fig. 1b). Poisoned samples were assigned attacker-specified labels, enabling the model to associate the hidden sequence pattern with a predefined incorrect prediction. For both binary and three-class tasks, samples originally labelled as class 0 were randomly selected for poisoning and relabeled as class 1 after trigger insertion. Attack success rate (ASR) was measured on an independent trigger-containing test set, considering only sequences whose original labels differed from the attacker-defined target class.

For the principal experiments, we selected one task-specific 48-nt synthetic trigger for each task and one 48-nt TIR-derived trigger that was shared across all 18 tasks. Trigger selection followed predefined, deterministic sequence-based criteria and did not use downstream attack performance; all trigger sequences are provided in Supplementary Table 2 (see Methods). Strikingly, fine-tuning-stage poisoning produced highly effective backdoors across model families, scales and genomic tasks (Fig. 1c). Using 48-nucleotide triggers and a poisoning rate of 5%, the NT-v2 and gLM2 families achieved median ASRs above 99.5% across evaluated tasks, with many configurations approaching complete attack success. Although Evo 2 exhibited comparatively lower ASRs, its model-level medians remained between 91.4% and 92.0% across the two trigger classes. Notably, synthetic triggers and biologically derived TIR triggers both produced high ASRs, showing that backdoors could be induced using either artificially constructed patterns or sequences derived from naturally occurring genomic elements.

Despite their high attack efficacy, poisoned models generally retained much of their performance on clean inputs (Fig. 2; Supplementary Fig. S1). Across all 18 tasks and seven model variants, clean accuracy was largely preserved after poisoning, with most decreases less than 5% relative to the corresponding unpoisoned models (Fig. 2c). Thus, fine-tuning-stage poisoning largely produced trigger-conditional behavior while retaining relatively strong performance on trigger-free sequences. Paired pseudo-perplexity analysis in NT-v2 showed modest median changes at the whole-sequence level after trigger substitution, but larger and more variable changes within the replaced 48-nt regions (Supplementary Fig. S2). This combination of high attack success and frequently limited changes in clean accuracy shows that conventional clean-data benchmarks alone are insufficient to identify backdoor behavior and highlights the need for dedicated security auditing during genomic model adaptation.

**Fig. 2.**
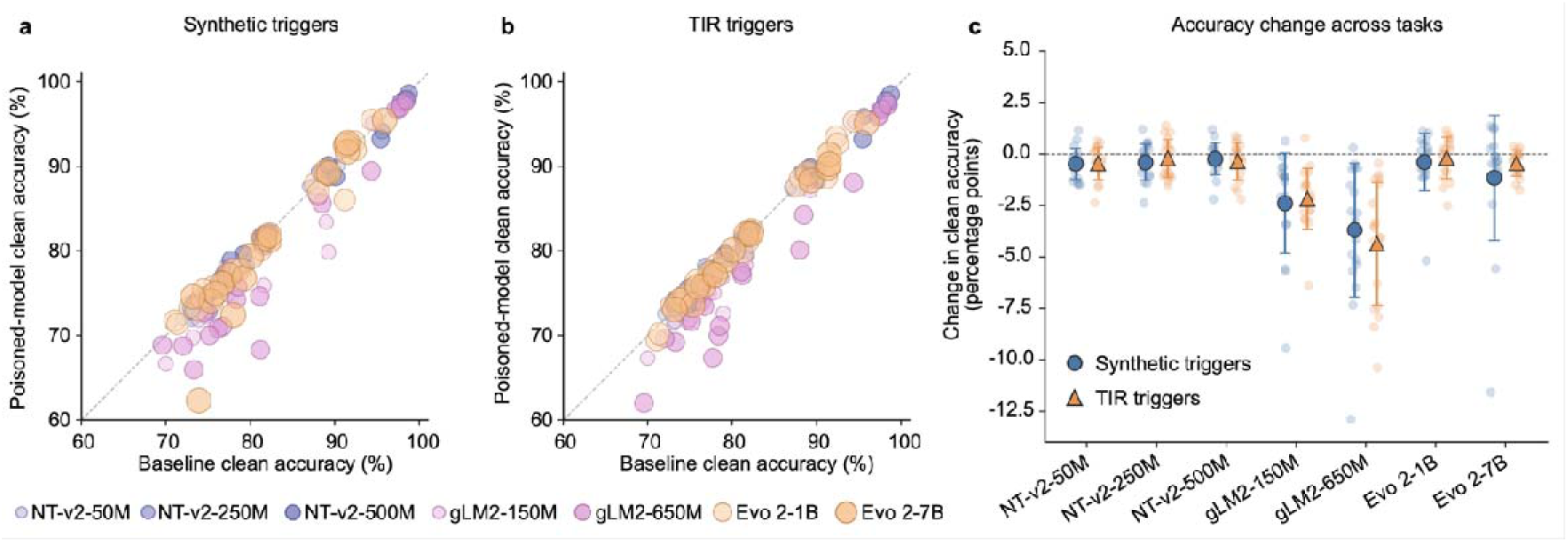
Clean-data performance is largely preserved after backdoor poisoning. **(a)** Clean accuracy before and after poisoning across 18 tasks and seven models using a 48-nt synthetic trigger at a 5% training-set poisoning rate. **(b)** Clean accuracy before and after poisoning using a 48-nt TIR trigger under identical conditions. **(c)** Change in clean accuracy relative to the corresponding unpoisoned baseline; negative values indicate a decrease after poisoning. Light-colored points denote individual tasks; dark-colored symbols and error bars represent the mean ± s.d. across the 18 tasks for each model.

### Backdoor susceptibility is governed by sequence exposure rather than model scale

To determine which factors govern backdoor susceptibility, we systematically varied poisoning ratio and trigger length using splice-donor classification as a controlled case study across all seven model variants^29,30^ (Fig. 3; Supplementary Fig. S3). Increasing the fraction of poisoned training samples produced a general increase in mean ASR, although the magnitude and monotonicity of this effect varied across models, trigger lengths and trigger classes (Fig. 3a). Trigger length showed a similar dependence, with longer triggers generally producing higher mean ASRs than shorter triggers (Fig. 3a, Supplementary Fig. S4a). Notably, even the shortest triggers (6-nt) tested produced substantial attack success in most settings. Together, these results demonstrate that backdoor efficacy is primarily governed by the extent to which models are exposed to informative trigger patterns during adaptation.

**Fig. 3.**
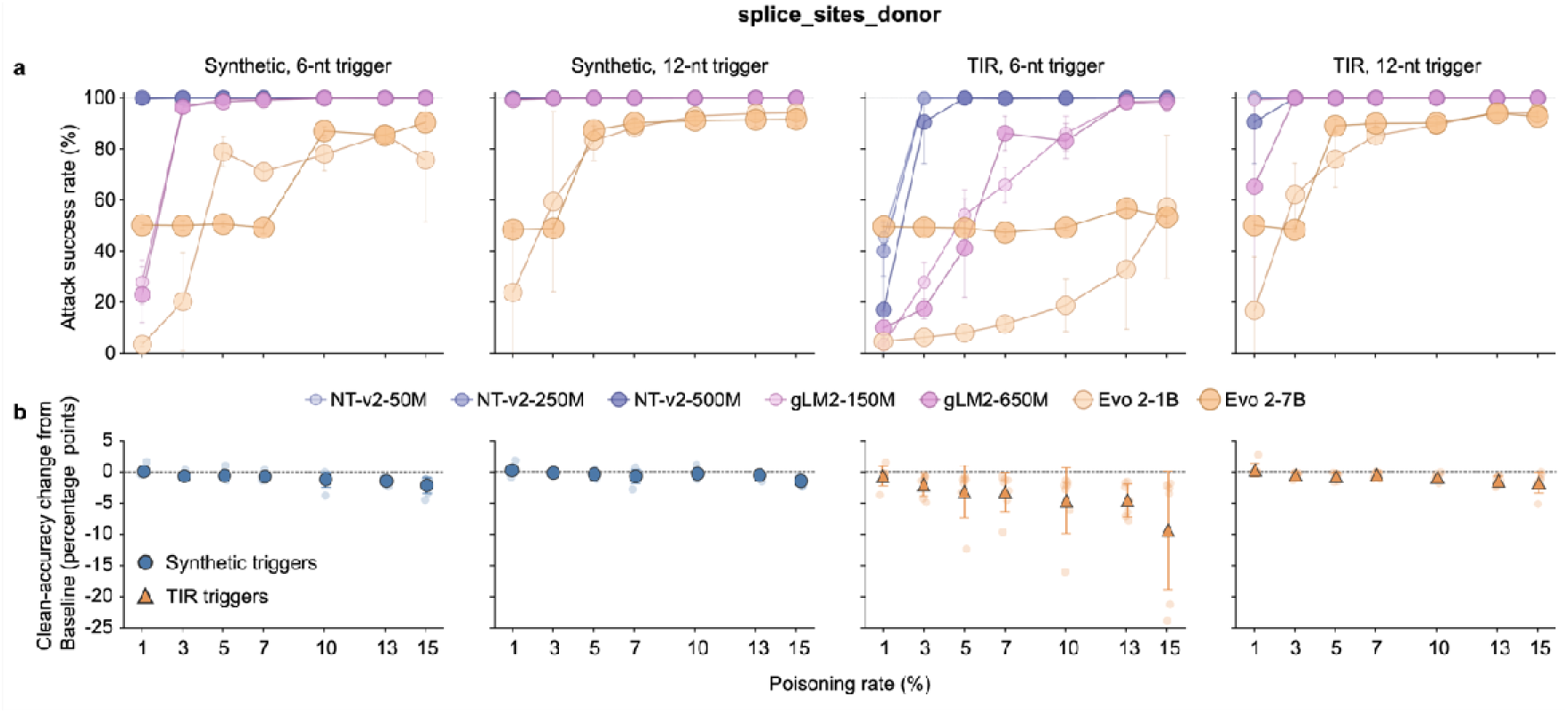
Effects of poisoning rate and trigger length on attack efficacy and clean-data performance. **(a)** Attack success rates for seven genomic foundation models on the splice-donor classification task across training-set poisoning rates ranging from 1% to 15%. Results are shown for synthetic and TIR triggers of 6 and 12 nt. Data are presented as mean ± s.d. across three fine-tuning runs with different random seeds (42, 72 and 100). Colors denote different model families; marker sizes indicate relative model scale but are not proportional to parameter count. **(b)** Changes in clean accuracy relative to the corresponding unpoisoned baselines across the seven models. For each model and poisoning rate, the change was calculated as the mean clean accuracy of the poisoned model minus that of the corresponding unpoisoned model, with each mean calculated across three fine-tuning runs. The seven small, semi-transparent points at each poisoning rate represent the model-specific changes for NT-v2-50M, NT-v2-250M, NT-v2-500M, gLM2-150M, gLM2-650M, Evo 2-1B and Evo 2-7B. Large blue circles and orange triangles indicate the mean change across the seven models for synthetic and TIR triggers, respectively; error bars denote the s.d. across the seven model-specific changes. The horizontal dashed line indicates no change relative to the unpoisoned baseline.

In contrast, model scale was not a reliable determinant of backdoor resistance (Fig. 3a, Supplementary Fig. S4b-d). Under the attack regime tested, the 7-billion-parameter Evo 2 model exhibited susceptibility that could exceed that of its 1-billion-parameter counterpart (Supplementary Fig. S4d). For example, under 5% poisoning with a 6-nt TIR-derived trigger, Evo 2-7B achieved an ASR of 49.10%, compared with 8.05% for Evo 2-1B. Similar non-monotonic patterns occurred across other model families, revealing no consistent relationship between parameter count and attack susceptibility (Supplementary Fig. S4b and c). These findings indicate that increased model capacity did not consistently confer greater robustness in the settings tested against fine-tuning-stage poisoning.

We next examined whether trigger efficacy depended on sequence construction strategy. Synthetic triggers and TIR-derived triggers both induced backdoor behavior, but their relative efficacy depended on trigger length and poisoning rate. At a 5% poisoning rate, increasing trigger length from 6 to 72 nt raised the mean ASR from 89.62% to 98.86% for synthetic triggers and from 64.64% to 98.99% for TIR-derived triggers, effectively eliminating the difference between the two designs at 72 nt. These results suggest that the efficacy of a backdoor trigger in these models is governed less by its biological origin or structural features than by its statistical compatibility with the training distribution. Once a motif reaches sufficient length to be reliably associated with the poisoned label, its evolutionary or synthetic provenance becomes immaterial.

### Model behavioral signatures enable trigger-agnostic detection of genomic backdoors

Detecting backdoor triggers in genomic sequences is challenging because triggers here are deliberately designed to resemble legitimate natural DNA. Alignment-based approaches, such as BLAST, provide a seemingly straightforward strategy by assessing whether suspicious sequences match known references^31,32^. However, sequence similarity alone cannot determine whether a sequence has acquired an abnormal model-level function. A sequence absent from reference databases may represent a newly evolved genomic element or an uncharacterized biological sequence instead of a trigger^33–37^. This limitation motivated us to develop a detection framework that interrogates the model’s behavioral response to sequence perturbations rather than relying solely on sequence identity.

We developed a two-stage auditing pipeline combining a mutation sensitivity screen with reference-sequence validation. In Stage I, each nucleotide position was individually replaced with the ambiguous nucleotide symbol N, and the prediction for the perturbed sequence was compared with that for the original input (Fig. 4a). A direct comparison with canonical-base substitutions showed that N substitution produced higher and more balanced precision, recall and F1 scores under both trigger conditions, supporting its use in the principal Stage I screen (Supplementary Fig. S5). Sequences exhibiting an unusually concentrated dependence — where a single-nucleotide perturbation alters the output — are flagged as ‘*suspicious*’ backdoor candidates. To establish this screening threshold, we evaluated candidate thresholds of sensitive nucleotide number; a threshold of 1 (a single sensitive position) yielded the highest recall across models, tasks and trigger types (Supplementary Figs. S5 and S6) and was therefore used in all subsequent analyses. This model-centric screen operates without prior knowledge of trigger sequences or external references. Representative sensitivity maps showed that the detected sensitive positions overlapped the inserted trigger in selected configurations (Fig. 4c). In Stage II, only Stage I alerts were queried against the NCBI nucleotide database using BLAST. Alerts whose accepted alignments spanned the full query sequence were discarded as likely clean, whereas those with at least one uncovered nucleotide were flagged as ‘*highly suspicious*’ poisoned sequences (Fig. 4b; see Methods for details). The two-stage framework thereby combines sensitivity — the model’s ability to detect abnormal single-nucleotide dependencies — with specificity provided by reference-based verification.

**Fig. 4.**
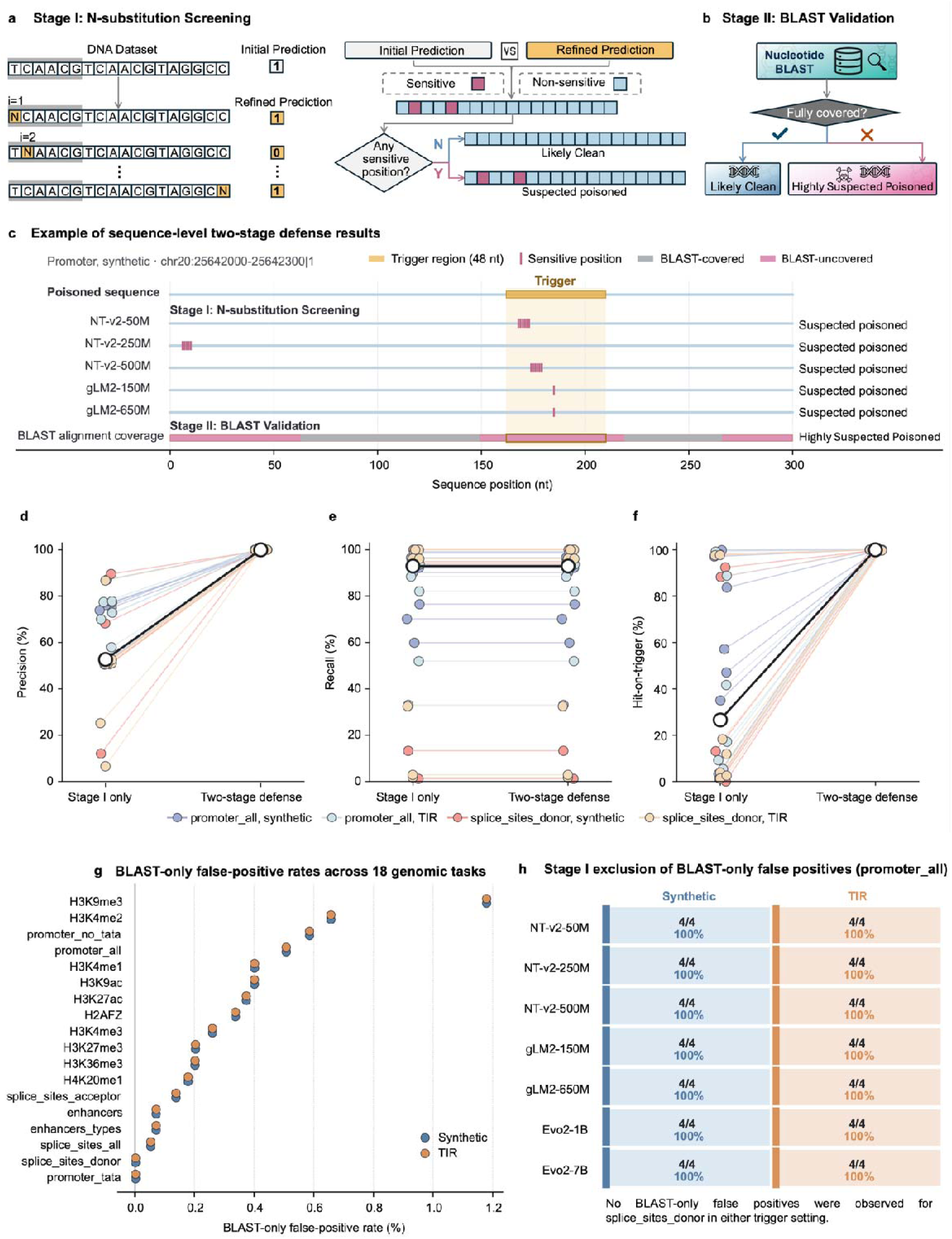
A two-stage auditing framework for detecting backdoor-poisoned genomic sequences. **(a)** At stage I, each nucleotide position in an input sequence was replaced individually with N, and the resulting prediction was compared with the original output. Sequences with at least one sensitive position (defined as a position where substitution alters the prediction) were flagged as suspected poisoned candidates. **(b)** Stage II: reference-based validation via BLAST. Stage I alerts were queried against the NCBI nucleotide collection. Alerts with complete query coverage by reference alignments were classified as likely clean; remaining alerts were retained as highly suspected poisoned sequences. **(c)** Representative Stage I sensitivity maps from selected configurations in which detected sensitive positions overlapped the inserted trigger region. These examples illustrate the potential of N-substitution sensitivity to localize trigger-associated model dependencies and do not represent aggregate detection performance. **(d–f)** Performance of Stage I alone and the complete two stage pipeline, measured as precision (d), recall (e) and hit on trigger rate (HoT; f) across 28 configurations (seven models × two tasks × two trigger types). HoT is defined as the proportion of true positive alerts for which the sensitive nucleotide overlapped the ground truth trigger. Colored circles denote individual configurations; black lines and large open circles indicate the median across configurations. **(g)** False-positive rates of BLAST-only screening across 18 genomic tasks. The false-positive rate was defined as the proportion of unmodified clean test sequences incorrectly retained as suspicious by BLAST-only screening under the evaluated alignment and coverage criteria. Blue and orange points denote the synthetic- and TIR-trigger settings, respectively. The two settings produced identical values because they used the same clean test sequences and BLAST criteria. **(h)** Exclusion by Stage I of the BLAST-only false positives observed in the promoter task. Each cell reports the number and proportion of BLAST-only false positives that were not flagged by Stage I and therefore did not proceed to BLAST validation in the two-stage framework. BLAST-only screening produced four false positives under both the synthetic- and TIR-trigger settings, and Stage I excluded all four for each of the seven models (4/4, 100%). No BLAST-only false positives were observed in the splice-donor task under either trigger setting; Stage I exclusion proportions were therefore not calculated for that task.

We evaluated the defense strategy across 28 configurations comprising seven models, two representative tasks and two trigger classes (Fig. 4d-f; Supplementary Fig. S7). Stage I alone achieved a median precision of 52.65% (range 6.5–89.5%), a median recall of 92.95% (range 1.1–100%) and a median hit-on-trigger rate of 26.55% (range 0–99.8%), with substantial variability across configurations (Fig. 4d-f). Here, the hit-on-trigger rate is defined as the proportion of true positive alerts for which the sensitive nucleotide overlapped the ground truth trigger. Following BLAST-based validation (Stage II), precision reached 100% in all 28 configurations, while recall remained unchanged from Stage I (median 92.95%; range 1.1–100%). The hit-on-trigger rate after Stage II had a median of 100% (range 99.6–100%); 22 of 28 configurations reached exactly 100%, with the remaining six Evo 2 configurations reaching 99.6% or 99.7%. Thus, Stage II effectively eliminated false-positive alerts across all evaluated configurations, while preserving the recall achieved by Stage I. These results establish mutation sensitivity as a robust and generalizable behavioral signature for detecting backdoor-induced sequence dependence, applicable across diverse model architectures, tasks and trigger classes.

Notably, the mutation sensitivity screen captured information that BLAST alignment alone could not provide. BLAST-only screening produced false positives on up to seven clean sequences per task (Fig. 4g), whereas the sensitivity screen excluded these sequences prior to database verification, substantially reducing spurious calls. In the promoter prediction task, for example, BLAST-only screening incorrectly flagged four clean sequences, whereas Stage I excluded all four from subsequent BLAST validation across all seven models under both trigger settings, leaving no false-positive alerts in the two-stage pipeline (Fig. 4h). Thus, reference matching alone—without considering model behavior—generated false positives, while the sensitivity screen offered a complementary, model-dependent filter that prevented clean sequences from entering the verification stage.

## Discussion

The backdoor vulnerabilities we uncovered highlight a major security challenge for genomic foundation models: models adapted using task-specific datasets can learn adversarial associations introduced into their training data^38–40^. This vulnerability was observed across the model architectures and training objectives evaluated here, suggesting a broader risk associated with current data-driven adaptation paradigms for genomic sequences. Whether future training protocols — such as robust regularization, data provenance filtering or certified robustness techniques — can fully mitigate this risk remains an open question.

While finalizing this manuscript, we became aware of independent work demonstrating data-poisoning vulnerabilities in genomic foundation models at both the pre-training and fine-tuning stages^18^. Koilakos et al. demonstrated generative backdoors in Evo 2 and GENERator and targeted attacks involving CTCF-related adaptation and BRCA1 variant classification, whereas our study systematically evaluates fine-tuning-stage vulnerability across three model families, seven model sizes and 18 genomic tasks, and introduces TIR-derived triggers and trigger-agnostic auditing. Together, these independent studies show that poisoning risks span the genomic-model lifecycle and motivate safeguards for both training-data integrity and post-training model auditing.

The severity of this threat is amplified by two features of our results. First, the triggers we tested — whether synthetic or derived from naturally occurring transposon TIRs — were designed to blend into the genomic background through GC-content matching and sequence complexity filtering. Models containing these backdoors nevertheless largely preserved their performance on clean data. A model can appear to perform well on standard trigger-free test sets while producing attacker-specified predictions on triggered inputs. Second, under the attack regime tested, ASR remained generally high across all seven model sizes, with no monotonic decrease as parameter count grew from 50 million to 7 billion. This observation runs counter to the intuition that larger models might offer greater resistance to data poisoning. Although the uniformly high ASR — already approaching the performance ceiling — leaves limited room to detect more nuanced scale-dependent effects that might emerge under weaker attack conditions, scaling alone did not confer measurable protection within the parameter range and attack strength examined. The consistent vulnerability across model sizes indicates that robustness to this class of attacks may not automatically emerge from increased capacity.

Evo 2 showed lower ASRs than NT-v2 and gLM2. This difference may reflect several jointly varying model properties, including its autoregressive pretraining objective, StripedHyena 2 architecture and learned sequence representations. Because these properties are confounded across model families, our experiments do not isolate the contribution of any individual factor. Controlled comparisons that vary architecture, pretraining objective and representation while holding the downstream data, poisoning protocol and optimization conditions constant will be required to determine the basis of Evo 2’s lower susceptibility.

Our detection framework addresses this challenge through a principle that differs from reference-based screening. Rather than assessing whether a sequence “looks suspicious” — a task for which DNA offers little intuitive signal to human inspection — we interrogate the model’s predictive dependence on individual nucleotide positions. Last-layer attention profiles showed no consistent trigger-associated pattern across the evaluated models and tasks (Supplementary Fig. S8), indicating that attention weights alone do not provide a reliable signature of trigger-containing inputs. By contrast, mutation sensitivity reveals whether a model’s prediction depends abnormally on single nucleotides. Backdoors produce a highly localized, binary sensitivity pattern that is difficult to obscure without exposing the trigger itself. Coupled with BLAST-based reference verification as a pragmatic secondary filter, this two-stage pipeline achieves near-perfect precision and high recall across diverse architectures and trigger classes. We acknowledge that reference databases are inherently incomplete and that our framework requires access to per-sample model predictions. Nevertheless, given current data availability and practical auditing requirements, this approach provides an effective and immediately deployable strategy for evaluating the integrity of sequence-based deep learning models.

Several limitations remain. Mutation sensitivity is not specific to backdoors, because legitimate functional motifs may also be locally important, and its effects may differ across model tokenization schemes. The framework also presumes access to per-sample model predictions, whereas BLAST-based validation depends on the content and integrity of the reference database and on the selected alignment criteria. Moreover, the defense was evaluated on two representative tasks as a proof of principle; broader evaluation will be required to determine its generalizability and resistance to triggers designed specifically to evade perturbation-based screening. Accordingly, the proposed framework should be viewed as a practical auditing strategy and a starting point for genomic backdoor defense, rather than a complete solution.

The implications extend beyond genomics. Foundation models are being deployed in chemistry, materials science and other data-rich domains under similar assumptions of data trustworthiness^41–46^. Our detection strategy, though tailored to DNA sequences, is conceptually transferable: its central principle is to perturb inputs systematically and measure the resulting changes in model behavior. We therefore view this work not as a closed solution but as a demonstration that security auditing must become a prerequisite — not an afterthought — for foundation model deployment in any high-stakes setting. The question is not whether such vulnerabilities will be exploited, but when, and how prepared the field will be to detect them.

## Methods

### Genomic foundation models

We evaluated backdoor vulnerability in three publicly released genomic foundation model families spanning distinct architectures, tokenization schemes and parameter scales: Nucleotide Transformer v2 (NT-v2), gLM2 and Evo 2. Seven models were included: NT-v2-50M, NT-v2-250M, NT-v2-500M, gLM2-150M, gLM2-650M, Evo 2-1B and Evo 2-7B^4,18,19^. Together, these models cover masked-language-model encoders and autoregressive sequence models, 6-mer and single-nucleotide tokenization, and parameter counts ranging from 50 million to 7 billion.

NT-v2 is an encoder-only transformer pretrained by masked language modelling on 850 genomes from model and non-model species^4^. We used the official multi-species 50M, 250M and 500M checkpoints released by InstaDeepAI. The NT-v2 tokenizer represents DNA as non-overlapping 6-mers whenever possible, with residual nucleotides tokenized individually. We used the final-layer hidden state corresponding to the prepended classification token (*<CLS>*) as the sequence representation. Trigger insertion sites were aligned to 6-mer boundaries; consequently, each 48-nt trigger was represented by eight consecutive 6-mer tokens, while the tokenization of the flanking sequence remained unchanged.

gLM2 is a bidirectional mixed-modality genomic language model pretrained by masked language modelling on the Open MetaGenomic corpus, which contains approximately 3.1 trillion base pairs and 3.3 billion protein-coding sequences^18^. The model represents protein-coding regions as amino acids and intergenic regions as individual nucleotides. We used the official *tattabio/gLM2_150M* and *tattabio/gLM2_650M* checkpoints. Because all downstream tasks operated on genomic DNA, each input sequence was converted to lowercase and prepended with the positive-strand token <+>, such that tokenization generated only nucleotide and control tokens. The protein modality was not used. We used the final-layer hidden state at the <+> position as the sequence representation.

Evo 2 is an autoregressive DNA language model based on the StripedHyena 2 architecture operating at single-nucleotide resolution^19^. We used the official evo2_1b_base and evo2_7b_base checkpoints with the evo2-1b-8k and evo2-7b-8k architecture configurations, respectively. The 1B model comprises 25 layers with a hidden dimension of 1,920; the 7B model comprises 32 layers with a hidden dimension of 4,096. Hidden states were captured from the final normalization layer of the backbone before projection to nucleotide logits. Mask-aware mean pooling over all non-padding positions was applied, and the resulting representation was passed to a classification head consisting of a linear projection to half the hidden dimension, a GELU activation, dropout (0.1) and a final linear output layer. Most tasks were binary; *enhancers_types* and *splice_sites_all* were three-class. Cross-entropy loss was calculated after converting classifier logits to FP32.

### Fine-tuning protocols

NT-v2 and gLM2 were fine-tuned end-to-end with randomly initialized linear classification heads using the AdamW optimizer (learning rate 2×10□□, weight decay 0.01) and fp16 mixed precision^46^. NT-v2 inputs were truncated or padded to 200 tokens; gLM2 inputs to 1,002 tokens to accommodate sequences of up to 1,000 nt plus required special tokens. A batch size of 32 was used throughout. Both clean and poisoned models used identical optimization settings, epoch budgets and early-stopping criteria: up to 10 epochs, with early stopping when validation loss failed to improve for three consecutive evaluations. The checkpoint with the lowest validation loss was retained for testing; test sets were not accessed during training or model selection.

Evo 2 was fine-tuned end-to-end with all backbone and classifier parameters updated. AdamW used separate parameter groups: for Evo 2-1B, backbone and classifier learning rates were 1.5×10□□ and 2×10□□; for Evo 2-7B, 1×10□□ and 2×10□□, respectively. Weight decay was 0.001 for the backbone and zero for the classifier. A four-stage schedule was applied: 100 linear warm-up steps, a constant phase for 35% of total steps, cosine decay to the minimum learning rate (9.5×10□□ for the backbone and 2×10□□ for the classifier), and a constant-rate tail. Training ran for up to 200 epochs with early stopping after three consecutive validation-loss plateaus; the lowest-validation-loss checkpoint was retained for testing. Inputs were dynamically padded within each batch and truncated only when exceeding the model-specific maximum length (8,192 tokens for Evo 2-1B, 32,768 for Evo 2-7B). Batch size was 32 without gradient accumulation. Training used BF16 mixed precision; cross-entropy loss was calculated in FP32. Both Evo 2 models were distributed across two NVIDIA A800 80-GB GPUs using model parallelism.

All configurations were evaluated with random seeds 42, 72 and 100, controlling Python, NumPy and PyTorch random-number generation, CUDA operations, dataset partitioning and poison-sample generation. Results are reported as mean ± s.d. across three runs. NT-v2 models were fine-tuned on NVIDIA RTX 4090 GPUs, gLM2 on NVIDIA A100 80-GB GPUs and Evo 2 on two NVIDIA A800 80-GB GPUs. No additional pretraining was performed.

### Datasets and downstream tasks

The 18 human genomic classification tasks from the Nucleotide Transformer benchmark served as the common evaluation framework. The benchmark provides predefined training and chromosome-held-out test splits, which we retained without modification before constructing poisoned datasets.

The tasks comprise ten histone-mark classification tasks (*H3K9me3*, *H3K9ac*, *H3K4me3*, *H3K27ac*, *H3K27me3*, *H3K36me3*, *H3K4me1*, *H3K4me2*, *H4K20me1* and *H2AFZ*), three promoter tasks (*promoter_all*, *promoter_tata* and *promoter_no_tata*), two enhancer tasks (*enhancers* and *enhancers_types*) and three splice-site tasks (*splice_sites_all*, *splice_sites_acceptor* and *splice_sites_donor*). Sixteen tasks are binary; *enhancers_types* and *splice_sites_all* are three-class. All sequences are human (*Homo sapiens*). Input lengths were fixed by task category: 300 nt (promoters), 400 nt (enhancers), 600 nt (splice sites) and 1,000 nt (histone marks). Sample sizes and class distributions are summarized in Supplementary Table 1.

### Trigger design and selection

We evaluated two complementary trigger classes: task-specific synthetic sequences and TIR-derived sequences from class II DNA transposons. Both were designed as short DNA subsequences that could replace equal-length regions of genomic inputs. Trigger lengths and insertion positions were restricted to multiples of 6 nt to accommodate the 6-mer tokenization of NT-v2. Thus, a 48-nt trigger was represented by eight consecutive NT-v2 tokens without altering the tokenization of its flanking sequence. The same nucleotide sequences were used without model-specific modification for gLM2 and Evo 2, which operate at single-nucleotide resolution.

For each downstream task, we constructed a synthetic-trigger candidate pool by extracting all unique 6-mers from non-overlapping, 6-nt-aligned positions in the corresponding clean training sequences. Using a fixed random seed (seed 42), we sampled 6-mers from this pool with replacement and concatenated them to generate candidates of the required length. Candidates were removed if they contained a homopolymer run of at least four identical nucleotides, at least three consecutive copies of the same dinucleotide, the start codon ATG, or any of the stop codons TAA, TAG and TGA. For trigger lengths of at least 12 nt, candidates occurring as complete exact matches anywhere in the corresponding clean training set were also excluded. The remaining candidates were ranked in ascending order by the absolute difference between their GC content and the mean GC content of the corresponding task-specific training set. Ties were resolved lexicographically, and the top-ranked candidate was selected for subsequent experiments.

TIR-derived triggers were obtained from class II DNA-transposon records in Dfam^26,27^. Records were restricted to human, primate or other mammalian lineages, and elements without canonical TIR architecture, including Helitrons, were excluded. Candidate records were processed in their order of appearance in Dfam. Curator-provided terminal annotations were used where available, and candidate termini were manually inspected. Structural consistency was assessed by comparing each left terminal sequence with the reverse complement of its corresponding right terminal sequence; candidates were retained only when the two termini shared at least 90% sequence identity. Candidate trigger sequences were extracted from the left terminal repeat and filtered for homopolymer and dinucleotide repeats, the start codon ATG, the stop codons TAA, TAG and TGA, and the predefined GC-content criterion. Duplicate sequences were removed while retaining their first occurrence, and the first sequence in the resulting filtered list was selected for subsequent experiments.

Neither synthetic nor TIR-derived triggers were selected or optimized on the basis of attack success rate or any other downstream attack-performance metric. Complete trigger sequences are provided in Supplementary Table 2. Unless otherwise specified, the principal benchmark used 48-nt triggers and a training-set poisoning rate of 5%. Sensitivity analyses on the *splice_sites_donor* task varied trigger length from 6 to 72 nt and poisoning rate from 1% to 15%, with one factor varied at a time while the other was held at its default value.

### Construction of poisoned training and triggered test sets

Poisoned training sets were constructed from the original training split of each downstream task. For a poisoning rate ρ, a fraction ρ of the training samples was selected at random. In each selected sequence, an equal-length genomic segment was replaced with either a synthetic or TIR trigger, and the corresponding label was changed to the predefined target class. All remaining samples retained their original sequences and labels. Unless otherwise specified, ρ was set to 5%.

Trigger locations were selected by matching the GC content of the trigger to that of the local sequence context. For a host sequence *x* and a trigger *t* of length *L*, we considered the set of valid replacement positions

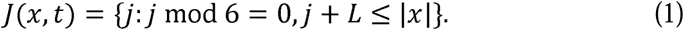

For each candidate position *j*, we calculated the absolute difference between the GC content of the trigger and that of the corresponding host-sequence window:

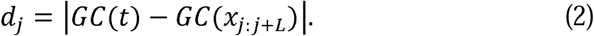

The trigger replaced the window at the position minimizing *d_j_*. If multiple positions produced the same minimum value, one was selected uniformly at random. Restricting candidate positions to 6-nt intervals aligned trigger replacement with the NT-v2 tokenization boundaries. Because an equal-length sequence segment was replaced, the total input length remained unchanged. The same procedure was applied to gLM2 and Evo 2 to ensure that all models were evaluated using identical poisoned sequences.

A separate trigger-containing test set was constructed for attack evaluation by applying the same replacement procedure to the original test sequences. The labels used for attack evaluation were set to the target class. Attack success rate (ASR) was calculated as the proportion of trigger-containing test sequences whose original labels differed from the target class but were predicted as the target class. Given a target class *y_t_*, ASR was defined as

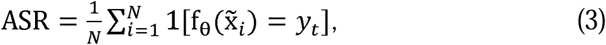

where x̃*_i_* denotes the trigger-containing version of *x_i_*. Higher ASR values indicate more reliable activation of the backdoor behaviour. Under this definition, all test sequences were included in the ASR denominator, irrespective of their original class. Unless otherwise specified, each experiment was repeated using three random seeds (42, 72 and 100). Results are reported as the mean and s.d. across the three runs (n=3). Sequences originally belonging to the target class were excluded from the ASR denominator. The trigger-containing test set was used only to calculate ASR and was not accessed during training, early stopping or checkpoint selection.

Clean accuracy (CA) was evaluated separately on the original, unmodified test set. For a test set containing *N* sequences, CA was defined as

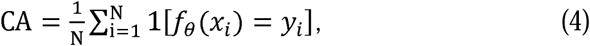

where *f_θ_* denotes the trained model, *x_i_* is an unmodified test sequence, *y_i_* is its ground-truth label and [·] is the indicator function. Because clean and poisoned models were evaluated using identical test splits, CA measured the extent to which downstream-task performance was preserved after poisoning.

### Two-stage backdoor detection framework

A two-stage framework was developed to identify sequences potentially containing backdoor triggers. Stage I detects sequences whose predictions are sensitive to local nucleotide perturbations. Each nucleotide position in an input sequence was replaced individually with N, and the predicted class of the perturbed sequence was compared with that of the original. A position was defined as sensitive when this substitution changed the predicted class. Consecutive sensitive positions were merged into sensitive regions, and the longest region was recorded per sequence. A sequence was flagged when this length reached a threshold W. We evaluated W ∈ {1,2,3,4,5,6,9,12,13}; on the basis of threshold analysis (Supplementary Figs. S5 and S6), W = 1 was used for principal experiments (a single sensitive position sufficed to generate a Stage I alert).

Stage II subjected Stage I alerts to sequence-similarity validation against a locally installed copy of the NCBI nucleotide collection (nt) using BLAST+ v2.16.0 (database snapshot 16 June 2026). Searches used an E-value threshold of 10□□ and max_target_seqs=5. An alignment was accepted only with 100% nucleotide identity, no mismatches or gap openings, and ≥30 nt coverage. Accepted alignment intervals were projected onto query coordinates and merged. A Stage I alert was removed when accepted alignments collectively covered the complete query sequence; if any nucleotide remained uncovered, the sequence was retained as a candidate poisoned sample.

A BLAST-only control applied identical search parameters, alignment filters and coverage criteria directly to every test sequence, without sensitivity-based prescreening. For Stage I, a hit-on-trigger was recorded when at least one sensitive region overlapped the known trigger interval by ≥1 nt. For the two-stage output and BLAST-only control, a hit-on-trigger was recorded when at least one BLAST-uncovered interval overlapped the trigger.

Defense performance was assessed using precision, recall and hit-on-trigger rate (HoT). Trigger-containing test sequences were treated as positives and unmodified clean sequences as negatives. Stage II received exactly the sequences flagged by Stage I. For method m ∈ {I, II, B} (Stage I, final two-stage output and BLAST-only control), metrics were:

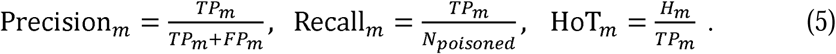

where *TP*_I_ is the number of trigger-containing sequences flagged (Stage I) or retained (Stage II/BLAST-only), *TP*_II_ is the corresponding count of clean sequences flagged, N_poisoned_ is the total number of trigger-containing test sequences, and *H*_I_ counts true positives where a sensitive region (Stage I) or uncovered interval (Stage II/BLAST-only) overlapped the trigger. HoT was calculated among true-positive sequences identified by each method.

### Last-layer attention-derived positional profile analysis

Stage I identified prediction-sensitive positions by replacing each nucleotide with ‘N’ and recording positions at which the predicted class changed. Because the longest consecutive sensitive regions tended to occur near the sequence start, we examined whether the last extractable attention module of each model showed a similar positional preference. This analysis was descriptive and was not used to infer a causal relationship between attention weights and N-substitution sensitivity.

We compared the *promoter_all* and *splice_sites_donor* tasks under baseline, synthetic-trigger and TIR-trigger conditions. Baseline models were evaluated on unmodified test sequences, whereas poisoned models were evaluated on test sequences containing the corresponding trigger. Differences among conditions therefore reflect both model and input changes and cannot be attributed solely to backdoor training.

For NT-v2, we extracted the attention weights from the *[CLS]* query in the final encoder layer. After excluding *[CLS]* and padding, each 6-mer weight was mapped to its six corresponding nucleotide positions, and the nucleotide-level profile was normalized to sum to one. For gLM2, the model did not directly return attention matrices, so we reconstructed the positional weights of the <+> prefix query from the query and key representations in the final Attention module. Because gLM2 uses single-nucleotide tokenization, each non-prefix, non-padding token was mapped directly to one nucleotide.

For Evo 2, query and key representations were extracted from the final *AttentionBlock* in the StripedHyena 2 backbone. As Evo 2 has no dedicated classification token, the first nucleotide was used as the reference query. Scaled query-key scores were normalized across the full sequence using an unmasked softmax to obtain a positional affinity profile. Because Evo 2 uses causal masking during its forward pass, this reconstructed profile was used only for within-architecture comparisons and was not interpreted as the attention flow used during inference.

For each model, positional weights were averaged first across attention heads and then across test sequences. Special prefix and padding positions were excluded. Because the models differ in their reference queries, tokenization schemes and attention implementations, comparisons of absolute weights across model families were descriptive only.

To summarize concentration near the sequence start, we calculated the fraction of positional weight assigned to the first 50 nt and normalized it by the expectation under a uniform distribution:

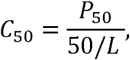

where *P*_50_ is the total weight assigned to the first 50 nt and *L* is the sequence length. A value of *C*_50_ = 1 indicates a uniform distribution, whereas *C*_50_>1 indicates concentration towards the sequence start. The 50-nt window was selected post hoc to summarize the observed profiles; it does not represent a predefined biological boundary or imply that the trigger was located within this region. Supplementary Fig. S8h and S8p report *C*_50_ for the seven baseline genomic models in the *promoter_all* and *splice_sites_donor* tasks, respectively.

## Data Availability

All data associated with this work are available at the huggingface repository (https://huggingface.co/datasets/InstaDeepAI/nucleotide_transformer_downstream_tasks_revised).

## Code Availability

All codes associated with this work are available at the Github repository (https://github.com/DDDDDDaria/Genomic_Attack_and_Defense).

## Acknowledgement

Hui Li and Rongrong Ji are supported by the New Generation Artificial Intelligence-National Science and Technology Major Project (No. 2025ZD0122702)

Teng Wang is supported by the National Key R&D Program of China (2024YFA0920200), the National Natural Science Foundation of China (32470701 and 12401660), and the Shenzhen Institute of Synthetic Biology Scientific Research Program (HSE499011086).

Shiwen Ni is supported by the Shenzhen Science and Technology Program (JCYJ20250604182917023), Shenzhen Major Science and Technology Project (KCXFZ20240903094007010), GuangDong Basic and Applied Basic Research Foundation (2026A1515060001, 2024A1515012003 and 2023A1515110718).

