## Supplementary Information for "Pervasive Backdoor Vulnerabilities in Genomic Foundation Models"

Shiwen Ni^1,*,^
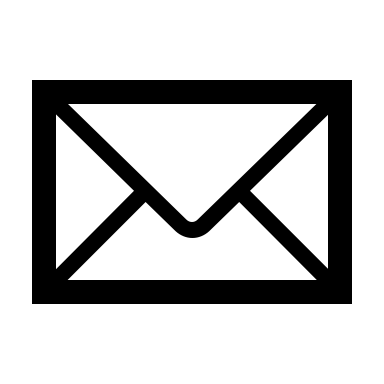
, Qianning Wang^1,*^, Chi Wei^2,*^, Xiaomin Ni^1^, Shuaimin Li^2^, Zixin Zhao^1^, Hui Li^3^, Rongrong Ji^3^, Ruifeng Xu^4,5^, Hamid Alinejad-Rokny^6^, Teng Wang^7,^
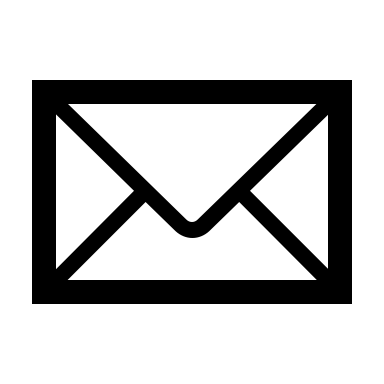
, Min Yang^1,2,7,^
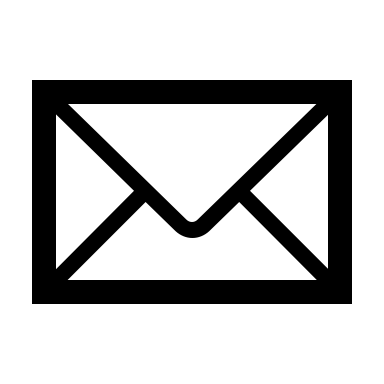

^1^Artificial Intelligence Research Institute, Shenzhen University of Advanced Technology, Shenzhen, China

^2^Shenzhen Key Laboratory for High Performance Data Mining, Shenzhen Institutes of Advanced Technology, Chinese Academy of Sciences, Shenzhen, China

^3^Key Laboratory of Multimedia Trusted Perception and Efficient Computing, Ministry of Education of China, Xiamen University, Xiamen , China

^4^College of artificial intelligence, Harbin Institute of Technology, Shenzhen, China

^5^Shenzhen loop area institute, Shenzhen, China

^6^School of Biomedical Engineering, UNSW Sydney, Sydney, Australia

^7^State Key Laboratory of Quantitative Synthetic Biology, Shenzhen Institute of Synthetic Biology, Shenzhen Institutes of Advanced Technology, Chinese Academy of Sciences, Shenzhen, China

^*^ These authors contributed equally to this work.

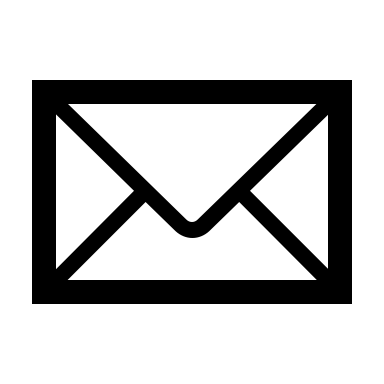

**Supplementary Figures**

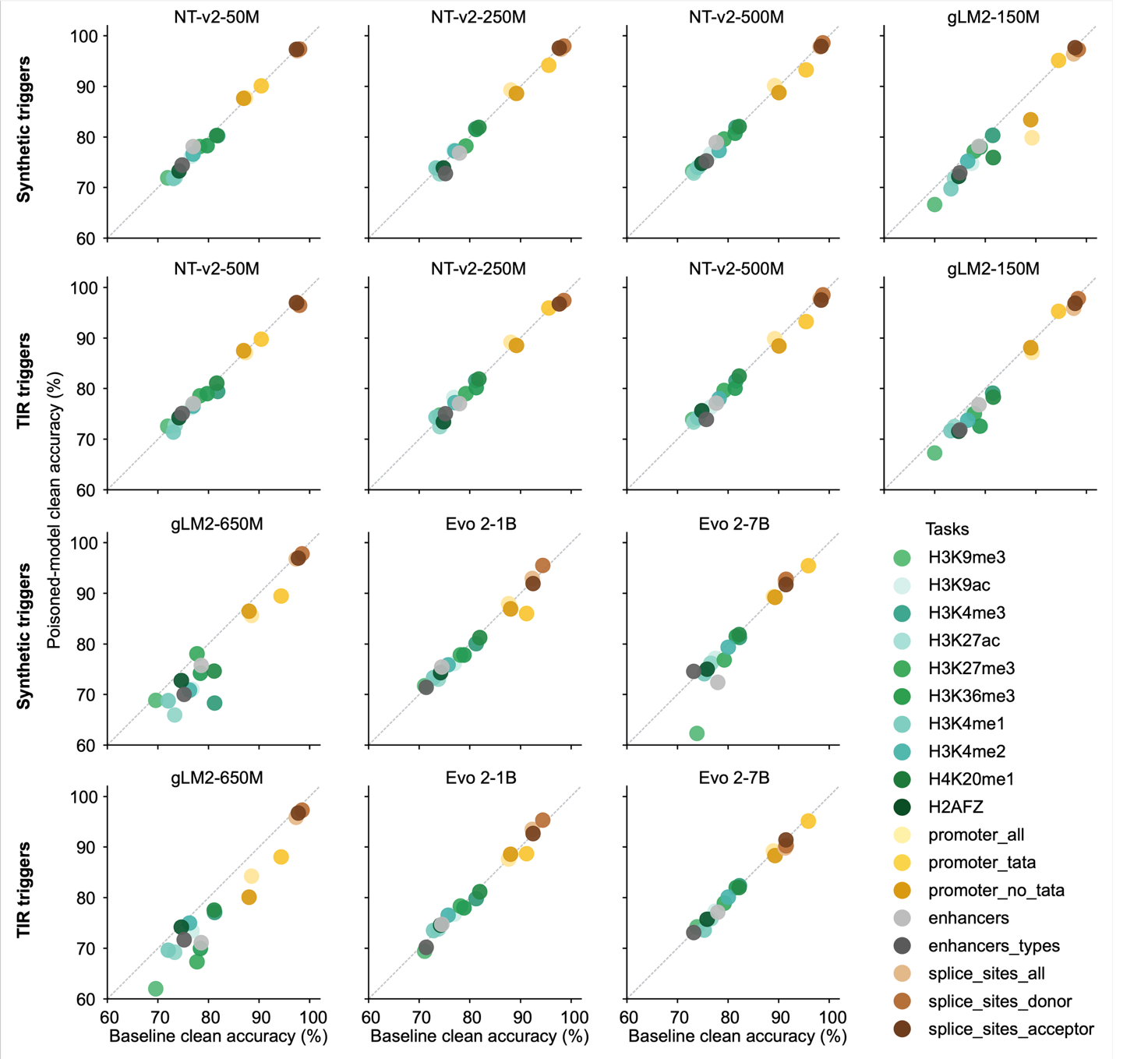

**Supplementary Figure S1 | Task-level clean accuracy of individual models before and after backdoor poisoning.** Clean accuracy of unpoisoned versus poisoned models for seven genomic foundation models across 18 downstream tasks, shown separately for synthetic and TIR triggers (48‑nt, 5% poisoning rate). Each point represents one task; colors indicate task identity. The dashed diagonal line denotes equal performance before and after poisoning. Values are means from three independent fine‑tuning runs. Clean accuracy was evaluated on unmodified, trigger‑free test sequences.

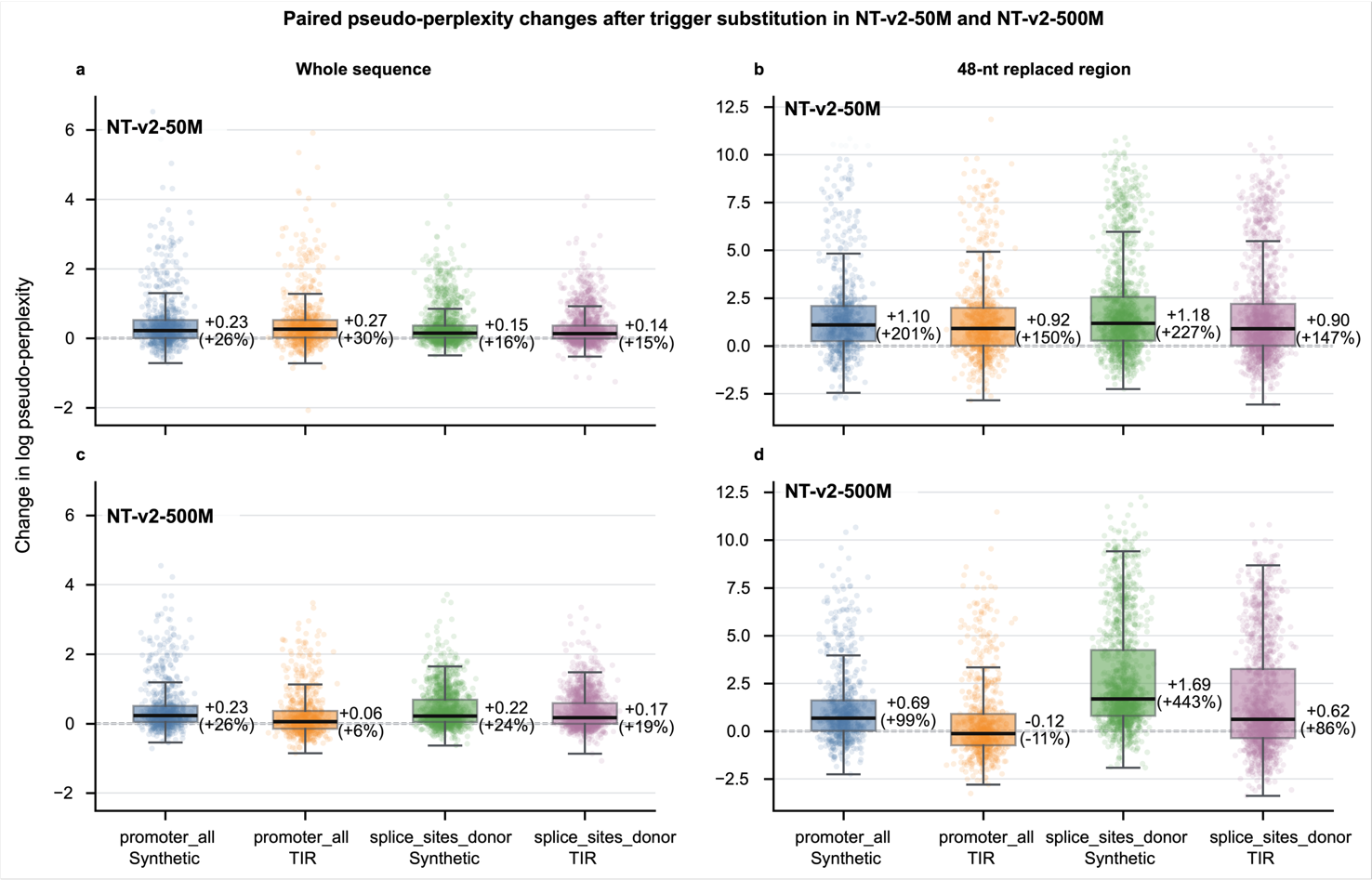
**Supplementary Figure S2 | Paired changes in pseudo-perplexity after trigger substitution in NT-v2 models.** Log pseudo-perplexity was compared between each unmodified test sequence and its trigger-containing counterpart for NT-v2-50M (a,b) and NT-v2-500M (c,d). Analyses were performed over the complete input sequence (a,c) or restricted to the 48-nt replaced region (b,d). Each point represents the paired difference for one test sequence, calculated as the log pseudo-perplexity of the trigger-containing sequence minus that of the corresponding unmodified sequence. Positive values indicate increased pseudo-perplexity after trigger substitution, whereas negative values indicate decreased pseudo-perplexity; horizontal dashed lines indicate no change. Blue, orange, green and purple denote promoter synthetic, promoter TIR-derived, splice-donor synthetic and splice-donor TIR-derived conditions, respectively. Each trigger condition included 792 paired promoter sequences or 1,520 paired splice-donor sequences. Boxes show the interquartile range, centre lines indicate the median, and whiskers extend to 1.5 times the interquartile range. The first value adjacent to each box indicates the median change in log pseudo-perplexity. The value in parentheses gives the corresponding percentage change in pseudo-perplexity, calculated as $100[\exp(\triangle)-1]$, where $\triangle$ is the median log pseudo-perplexity difference. For example, a median difference of (+0.23) corresponds to an approximately 26% increase in pseudo-perplexity.

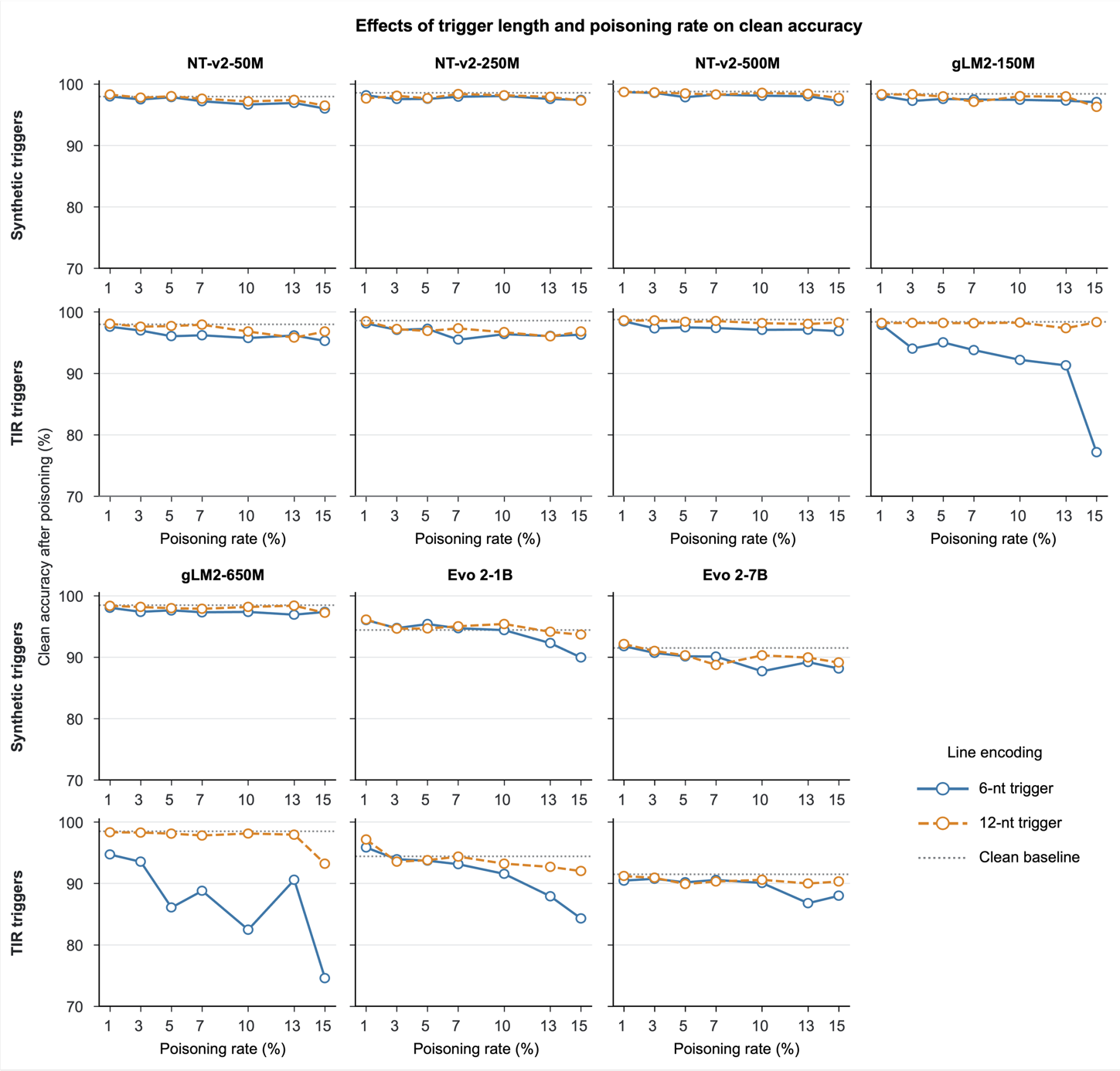

**Supplementary Figure S3 | Clean accuracy across poisoning rates and trigger lengths on the splice_sites_donor classification task.** Clean accuracy of unpoisoned versus poisoned models for seven genomic foundation models, shown separately for synthetic and TIR triggers. Poisoning rates range from 1% to 15%; solid and dashed lines denote trigger lengths of 6 and 12 nt, respectively. Values are means from three independent fine‑tuning runs. Clean accuracy was evaluated on unmodified, trigger‑free test sequences. The horizontal dashed line indicates no change relative to the unpoisoned baseline.

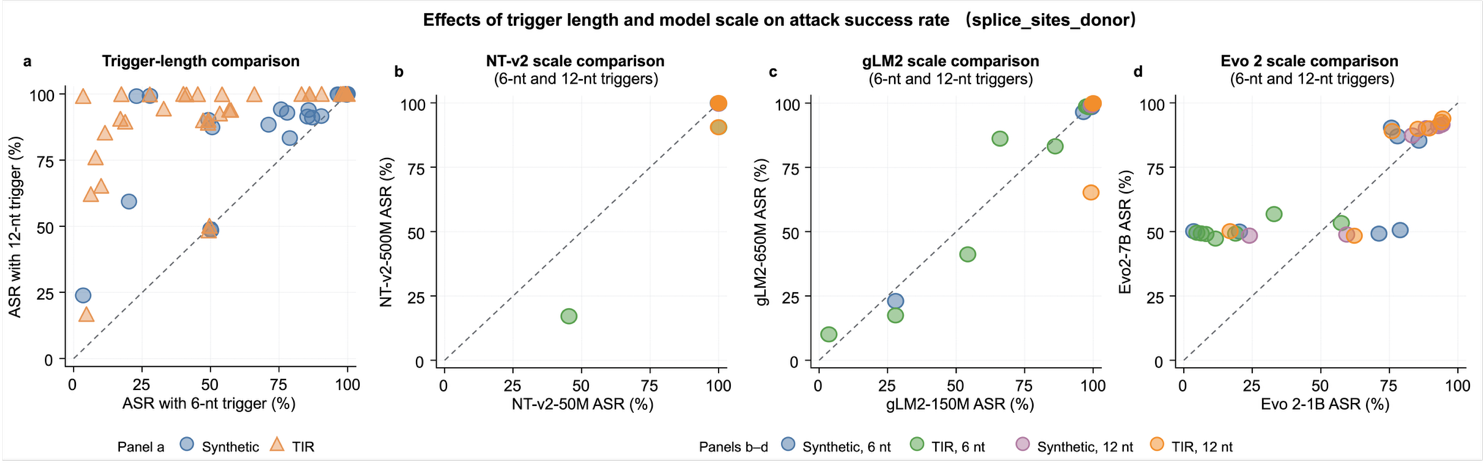

Supplementary Figure S4 | Effects of trigger length and model scale on attack success in the splice_sites_donor task. **(**a**)** Paired comparison of attack success rates (ASRs) obtained with 6-nt and 12-nt triggers under matched model, trigger-type and poisoning-rate conditions. Each point represents the mean ASR from three fine-tuning runs for one matched configuration. Blue circles and orange triangles denote synthetic and TIR-derived triggers, respectively. **(**b**-**d**)** Paired comparisons of ASR between NT-v2-50M and NT-v2-500M (b), gLM2-150M and gLM2-650M (c), and Evo 2-1B and Evo 2-7B (d) under matched trigger-length, trigger-type and poisoning-rate conditions. Blue, green, purple and orange circles denote synthetic 6-nt, TIR-derived 6-nt, synthetic 12-nt and TIR-derived 12-nt triggers, respectively. In a, points above and below the diagonal indicate higher and lower ASR, respectively, for the 12-nt trigger relative to the matched 6-nt trigger. In b–d, points above and below the diagonal indicate higher and lower ASR, respectively, for the larger model relative to the smaller model from the same family. Twelve-nucleotide triggers more frequently produced equal or higher ASRs than their matched 6-nt counterparts, whereas the direction and magnitude of the model-size effect varied across model families and attack conditions.

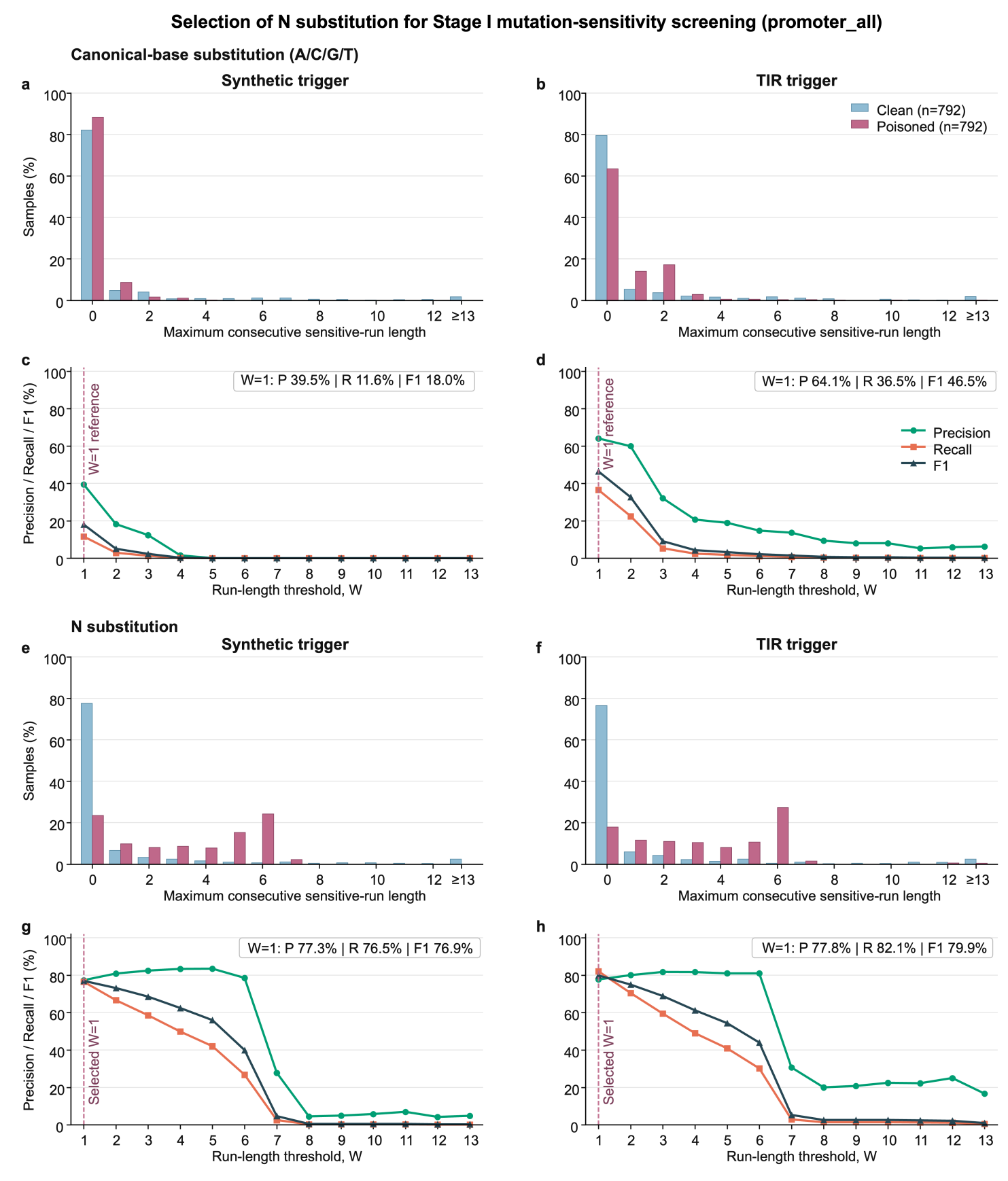

**Supplementary Figure S5 | Comparison of canonical-base and N substitutions for Stage I sensitivity screening.** Canonical-base and N-substitution screening were compared using NT-v2-500M on the *promoter_all* task under 48-nt synthetic- and TIR-trigger conditions. Each condition included 792 trigger-containing source-class sequences and their 792 corresponding unmodified sequences. For canonical-base screening, each nucleotide was individually replaced with each of the other three canonical bases, and a position was defined as sensitive if at least one substitution changed the predicted class. For N-substitution screening, each nucleotide was replaced with *N*, and a position was defined as sensitive if this replacement changed the predicted class. **(a, b)** Distributions of the maximum consecutive sensitive-run length in clean and trigger-containing sequences under canonical-base substitution for synthetic (a) and TIR-derived (b) triggers. **(c, d)** Precision, recall and F1 score of canonical-base screening across run-length thresholds $W$ for synthetic (c) and TIR-derived (d) triggers. **(e,f )** Corresponding distributions under N substitution. **(g, h)** Precision, recall and F1 score of N-substitution screening across thresholds for synthetic (g) and TIR-derived (h) triggers. Poisoned and clean sequences were treated as positive and negative samples, respectively. A sequence was flagged when its maximum consecutive sensitive-run length was at least $W$; at $W=1$, a single sensitive position was sufficient, without requiring adjacent sensitive positions. At $W=1$, the F1 scores for canonical-base and N-substitution screening were 18.0% and 76.9%, respectively, under the synthetic-trigger condition, and 46.5% and 79.9% under the TIR-trigger condition. N substitution produced more detectable prediction changes in trigger-containing sequences and a more balanced precision–recall profile, supporting its use in the principal Stage I screen.

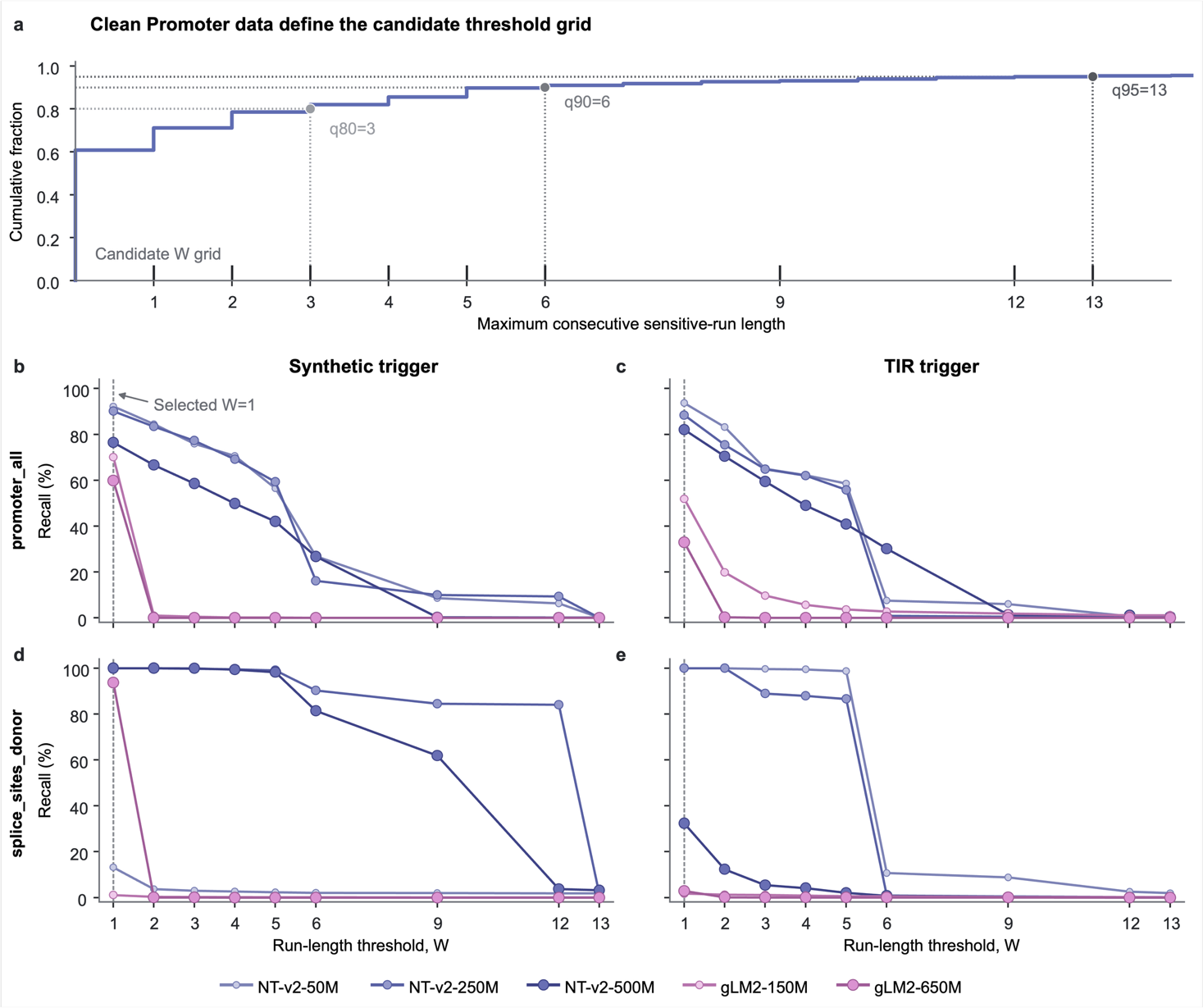

**Supplementary Figure S6 | Threshold selection for Stage I mutation‑sensitivity screening.**

**(a**) Empirical cumulative distribution of the maximum consecutive sensitive-run length in clean promoter sequences. Results were obtained using the clean NT-v2-50M model fine-tuned with seed 42 on the promoter_all task and evaluated on 1,584 unmodified test sequences of 300 nt. Each nucleotide was individually replaced with $N$, and a position was defined as sensitive when the substitution changed the predicted class relative to the original sequence. For each sequence, the maximum number of consecutive sensitive positions was recorded as *R*. The cumulative fraction at *r* was calculated as the proportion of clean sequences satisfying $R\leq r$. The 80th, 90th and 95th percentiles occurred at $R=3$, (6) and (13), respectively. These percentiles, together with intermediate values, informed the candidate threshold grid $W\in\{1,2,3,4,5,6,9,12,13\}$.

**(b–e**) Recall of Stage I across candidate thresholds for synthetic and TIR triggers in the promoter_all (b,c) and splice_sites_donor (d,e) tasks. Recall was defined as the proportion of trigger-containing source-class test sequences whose maximum consecutive sensitive-run length was at least $W$. Curves show results for poisoned NT-v2-50M, NT-v2-250M, NT-v2-500M, gLM2-150M and gLM2-650M models. Recall was non-increasing as $W$ increased, but its absolute value and rate of decline varied substantially among models, tasks and trigger classes. At $W=1$, recall ranged from 1.12% to 100% across the evaluated configurations. Because $W=1$ yielded the highest or joint-highest recall in every configuration, it was selected as a recall-prioritizing threshold for the principal Stage I screen (vertical dashed line). This selection represents a permissive first-stage screening rule and should not be interpreted as a globally optimal trade-off between false-positive rate and recall.

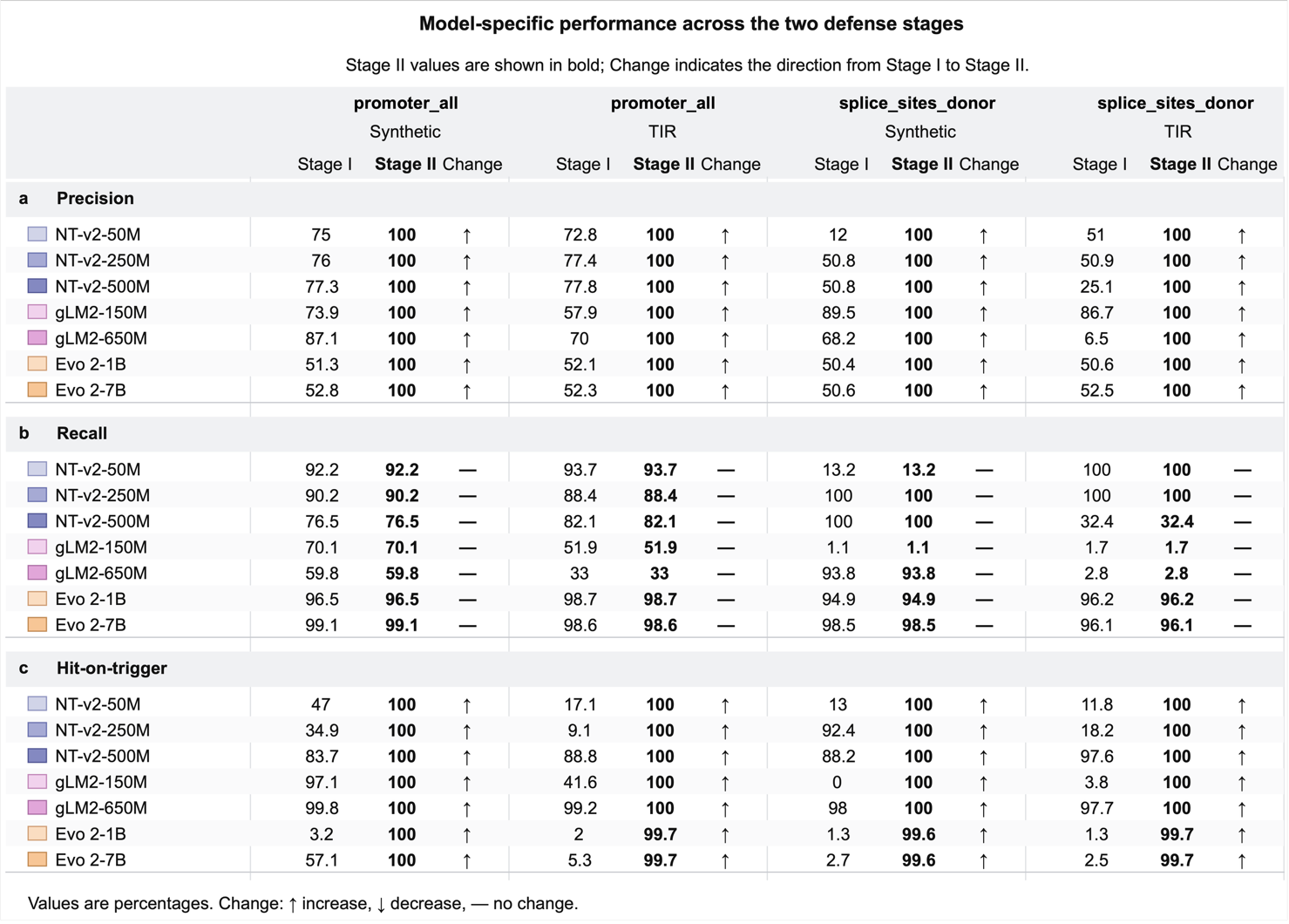

Supplementary Fig**ure** S7 | Model-specific performance of the two-stage auditing framework. Model-specific precision (a), recall (b) and hit-on-trigger rate (HoT; c) are shown for seven genomic foundation models evaluated on the promoter_all and splice_sites_donor tasks under synthetic- and TIR-trigger conditions. Stage I denotes N-substitution-sensitivity screening, and Stage II denotes the final output after BLAST-based reference validation. Rows correspond to individual models, and columns correspond to the four task-trigger combinations. Values are percentages. Stage I and Stage II values are reported separately, with Stage II values shown in bold. In the Change columns, “↑”, “↓” and “—” indicate an increase, decrease or no change, respectively, from Stage I to Stage II. Colored swatches identify the seven model variants, with related colors denoting models from the same family. For Stage I, HoT was defined as the proportion of true-positive alerts in which at least one sensitive region overlapped the inserted trigger. For Stage II, HoT was defined as the proportion of true-positive alerts in which at least one BLAST-uncovered region overlapped the trigger.

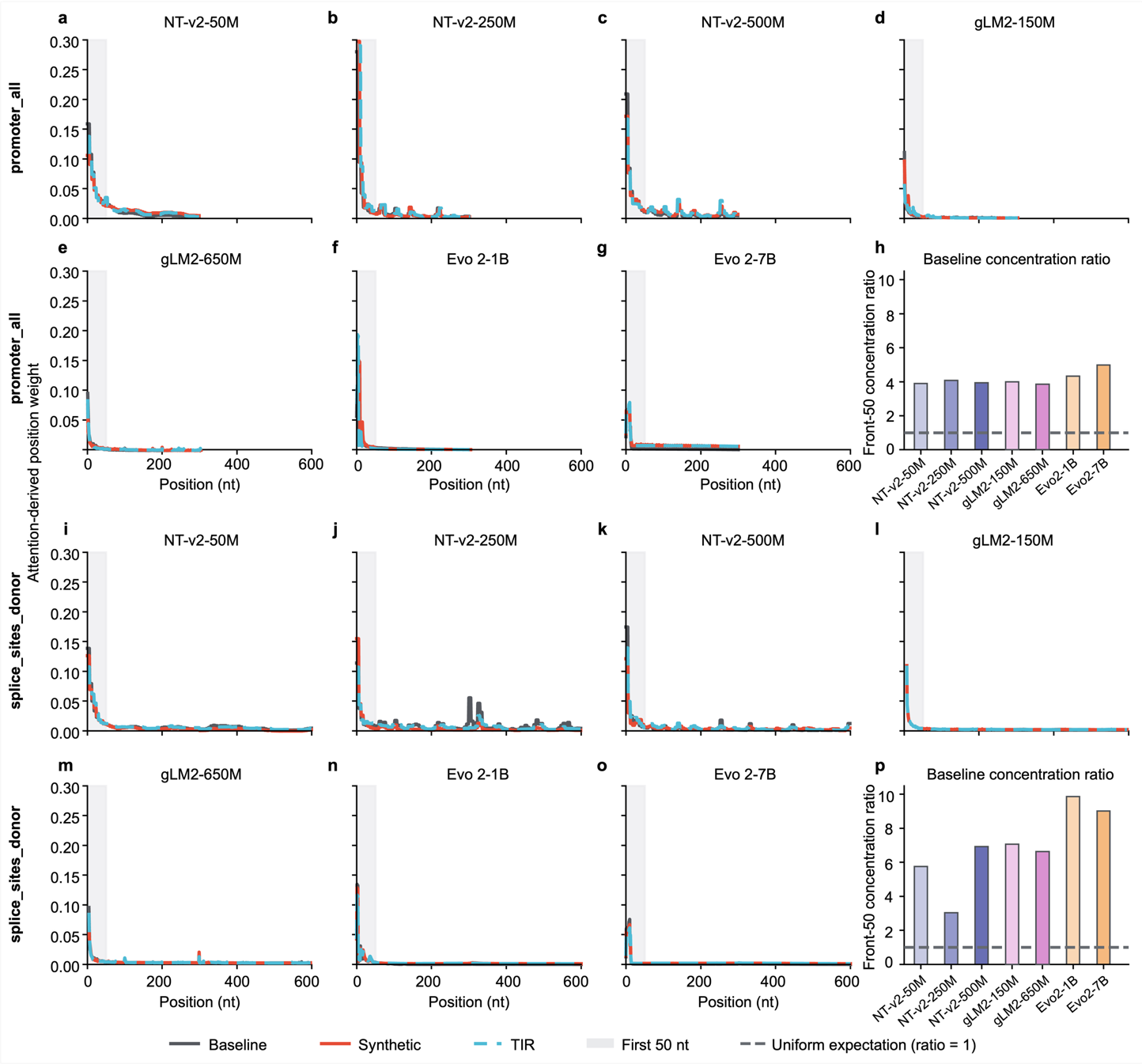

**Supplementary Figure S8 | Last-layer attention profiles do not reveal a consistent trigger-associated pattern across genomic foundation models.**

**(a–g)** Last‑layer attention profiles of seven genomic foundation models evaluated on the promoter task under baseline, synthetic-trigger and TIR-trigger conditions.

**(h)** Baseline attention concentration within the first 50 nucleotide positions for the promoter task.

**(i–o)** Corresponding attention profiles on the splice‑donor prediction task.

**(p)** Baseline attention enrichment within the first 50 positions for splice‑donor prediction. Grey, red and blue curves denote baseline, synthetic‑trigger and TIR‑trigger conditions, respectively; shading indicates the first 50 positions. Enrichment is defined as the fraction of attention assigned to the first 50 positions divided by the fraction expected under uniform attention (50/L, where L is the sequence length); a ratio of 1 indicates uniform attention, and a ratio greater than 1 indicates preferential attention to the sequence start. The horizontal dashed line in (**h)** and (**p)** marks a ratio of 1. Baseline attention was concentrated near the sequence start across the evaluated genomic models, particularly for the splice-donor task, whereas trigger-associated changes varied among models and showed no consistent pattern.

**Supplementary Table 1 | Summary of the 18 classification tasks used in this study.**

| Task | Category | Classes | Training sequences | Test sequences | Sequence length  (nt) |
| --- | --- | --- | --- | --- | --- |
| H3K9me3 | Histone mark | 2 | 27,438 | 850 | 1,000 |
| H3K9ac |  |  | 23,274 | 1,004 |  |
| H3K4me3 |  |  | 17,468 | 776 |  |
| H3K27ac |  |  | 30,000 | 1,616 |  |
| H3K27me3 |  |  |  | 3,000 |  |
| H3K36me3 |  |  |  | 3,000 |  |
| H3K4me1 |  |  |  | 3,000 |  |
| H3K4me2 |  |  |  | 2,138 |  |
| H4K20me1 |  |  |  | 2,270 |  |
| H2AFZ |  |  |  | 3,000 |  |
| promoter_all | Promoter | 2 | 30,000 | 1,584 | 300 |
| promoter_tata |  |  | 5,062 | 212 |  |
| promoter_no_tata |  |  | 30,000 | 1,372 |  |
| enhancers | Enhancer | 2 | 30,000 | 3,000 | 400 |
| enhancers_types |  | 3 |  |  |  |
| splice_sites_all | Splice site | 3 | 30,000 | 3,000 | 600 |
| splice_sites_acceptor |  | 2 |  |  |  |
| splice_sites_donor |  |  |  |  |  |

Supplementary Table 2 | **All t**rigger sequences used in this study. Synthetic triggers were generated separately for each task, except that enhancers and enhancers_types used the same 48-nt trigger. The same 48-nt TIR-derived trigger was used across all 18 tasks in the principal benchmark. Trigger lengths of 6, 12, 24, 36, 48, 60 and 72 nt were evaluated on the splice_sites_donor task.

| Trigger type | Task scope | Trigger length (nt) | Trigger sequence |
| --- | --- | --- | --- |
| Synthetic | H2AFZ | 48 | AGATTGGTGCGGAAACAAATATACAAGGGATTACCGATCGACTGGTCA |
| Synthetic | H3K27ac | 48 | CTCAGCTTGGACACTTCAGTCTATCGTTCTTCTTTACCAGAATCCACT |
| Synthetic | H3K27me3 | 48 | CGGAAGTCAAAGGTTGGATATACTATCAGACTACCAAGACGTTGCTAC |
| Synthetic | H3K36me3 | 48 | CTTACGAACTTCAATTTGTACGTCTTCCCACAATACATCGAGACCGGA |
| Synthetic | H3K4me1 | 48 | ACGTGTATACCGGTTGGATATACTATCAGACTACCAAGACGTTGGTTC |
| Synthetic | H3K4me2 | 48 | GCTGTTTGGCTACCAAGATCCTACAGCCACGAAGTACAAATATTGCTA |
| Synthetic | H3K4me3 | 48 | AGATACACTTCAACCCGAGCCAAATATCCGTTGCCGAAATTTGCGTCT |
| Synthetic | H3K9ac | 48 | CTCCGTCCGGTGTATCTCGGCATCGATTATTATTCAGAAGTTTCGATA |
| Synthetic | H3K9me3 | 48 | ATCCTCCGAGGATATTACTTCTGGTTGTCGCAGGATTTATATCAGAAC |
| Synthetic | H4K20me1 | 48 | TTCAATACGCCATCAAGGGCGTGTTTCAAGTTCTTCCTTGGTTACCAC |
| Synthetic | enhancers | 48 | TGCTACAATACGTGGGAGCTTACCAATTTCAATATCGTGCTTCTTGGA |
| Synthetic | enhancers_types | 48 | TGCTACAATACGTGGGAGCTTACCAATTTCAATATCGTGCTTCTTGGA |
| Synthetic | promoter_all | 48 | GACTTTCCGCTCAACGGCTGGTGTTCAAAGGCATTCGGAAGTTTCAGA |
| Synthetic | promoter_no_tata | 48 | AGCCGCACTCTGGGTATTCTTGGGCTCAGATCACTATCTCTGTTGTAC |
| Synthetic | promoter_tata | 48 | CCAACAATCACAACGACGCGGAGTCACCGTGGATATCATTCAACAGTT |
| Synthetic | splice_sites_acceptor | 48 | TGCCTTGCAAGAGCCATATTCACCAACTTATACAAGGTTGGCAACTTA |
| Synthetic | splice_sites_all | 48 | TTTGCCAGAGGGATTCCGATTCACACGTCAAGAGTTATACTATTCCAG |
| Synthetic | splice_sites_donor | 6 | TCAACG |
| Synthetic | splice_sites_donor | 12 | GATACGGACTTA |
| Synthetic | splice_sites_donor | 24 | TGCAGCTCATTTCCCTCAACAATT |
| Synthetic | splice_sites_donor | 36 | AAGGCCCAGAACCACAATCCTTATCATCTTATTCCA |
| Synthetic | splice_sites_donor | 48 | AGTTACTCATTGGGTCGCGTTTACTATTTCGTTCGTGCGATTCTCTTA |
| Synthetic | splice_sites_donor | 60 | AAACATTATTATATCAAACCGGTGTATTGGTGCTCCTCCTTTCAGAAAGCCGAGCCTCGT |
| Synthetic | splice_sites_donor | 72 | TCAACCTTGGTCAATCCATACGACGAAACACTATTATCGATCTATATCGACTATTCCTCAGCCAACGACCTT |
| TIR-derived | splice_sites_donor | 6 | CAGTCA |
| TIR-derived | splice_sites_donor | 12 | CAGTGGTCTTCA |
| TIR-derived | splice_sites_donor | 24 | CAGTAAGTCCTCACTTAACGTCGT |
| TIR-derived | splice_sites_donor | 36 | GGGTTTGGATCATAATCCCGAAAGACACAATCCCGA |
| TIR-derived | All 18 tasks | 48 | GGGTTTGGATCATAATCCCGAAAGACACAATCCCGAACGCCATAATCC |
| TIR-derived | splice_sites_donor | 60 | CAGTAAGTCCTCACTTAACGTCGTCGATAGGTTCTTGGAAACTGCGACTTTAAGCGAAAC |
| TIR-derived | splice_sites_donor | 72 | CAGTAAGTCCTCACTTAACGTCGTCGATAGGTTCTTGGAAACTGCGACTTTAAGCGAAACGACGTACTGTAT |
